# Small molecule-directed differentiation of submerged-cultured human nasal airway epithelia for respiratory disease modelling

**DOI:** 10.1101/2024.10.17.618953

**Authors:** Henriette H.M. Dreyer, Georgia-Nefeli Ithakisiou, Sacha Spelier, Malina K. Iwanski, Eugene Katrukha, Jonne Terstappen, Lisa W. Rodenburg, Loes A. den Hertog-Oosterhoff, Shannon M.A. Smits, Isabelle S. van der Windt, Lotte T. Azink, Linda H.M. Bijlard, Koen Passier, Sam F.B. van Beuningen, Robert Jan Lebbink, Eric G. Haarman, Cornelis K. van der Ent, Lukas C. Kapitein, Louis J. Bont, Jeffrey M. Beekman, Gimano D. Amatngalim

## Abstract

Submerged cultures of undifferentiated or transformed epithelial cells are widely used in respiratory research due to their ease of use and scalability. However, these systems fail to capture the cellular diversity of the human airway epithelium. In this study, we developed an *in vitro* model where cryopreserved human nasal epithelial cells, collected by brushings, are differentiated under submerged conditions on standard plastic cultureware. By applying small-molecule inhibitors targeting Notch and BMP signaling, we achieved efficient differentiation of cultures containing basal, secretory, and ciliated cells. This approach supports scalable culturing of both 2D epithelial monolayers and 3D organoids, validated as (personalized) disease models for primary ciliary dyskinesia, cystic fibrosis, and respiratory syncytial virus infection. This model offers a cost-effective, scalable platform that combines the simplicity of traditional cultures with the cellular complexity of the human airway epithelium, providing a valuable tool for respiratory disease research.

## Introduction

Despite advancements in sophisticated epithelial models, such as transwell-differentiated epithelia, organoids, and organ-on-a-chip systems, cell cultures on conventional plastic substrates under fluid-submerged conditions remain a cornerstone of biomedical research and drug development ^1^. Their widespread use is due to their ease of handling, scalability, and cost-effectiveness, making them ideal for generating large datasets and performing drug screening assays. However, these traditional models often lack the cellular diversity of native epithelia, leading to suboptimal experimental outcomes and contributing to higher drug development failure rates.

This limitation is particularly pronounced in respiratory research, where accurate *in vitro* models of the human airway epithelium are essential for successful drug discovery ^2^. For nearly 40 years, the air-liquid interface (ALI) culture model has been the gold standard for studying airway epithelial cells ^3^. In this model, primary airway basal progenitor cells are differentiated into secretory and ciliated cells under air-exposed conditions, generating monolayers that closely resemble the native airway epithelium ^4^. While highly valuable for research and drug validation, application of ALI-cultures is limited by the need for specialized transwell cultureware, which is costly and incompatible with certain assays. As a result, submerged cultures with undifferentiated airway basal cells and epithelial cell lines (e.g., 16HBE, A549, Calu-3) remain commonly used, despite their inability to replicate the complex cellular composition of ALI-cultures ^5^.

Given the need for accessible and biologically relevant airway models, nasal cells have emerged as a convenient and reliable source of human airway epithelial cells for *in vitro* studies ^6^. Nasal cells are easily collected through minimally invasive nasal brushing and share functional characteristics with bronchial epithelia, making them a valuable surrogate in respiratory research ^7^. Due to these advantages, nasal cultures are increasingly used in personalized disease models, especially for monogenic diseases such as primary ciliary dyskinesia (PCD) and cystic fibrosis (CF) ^6,8^. Additionally, nasal epithelial cells are widely applied to study respiratory virus infections, including respiratory syncytial virus (RSV) ^9,10^.

In this study, we aimed to bridge the gap between traditional submerged cultures and more advanced models by developing a simplified method in which cryopreserved human nasal airway epithelial cells are differentiated under submerged conditions on conventional culture plastics. By targeting Notch and BMP signaling pathways with small molecule inhibitors, we generated submerged cultures that more accurately reflect the composition of human airway epithelium, containing a mix of basal, secretory, and ciliated cells. This method enables the generation of 2D epithelial monolayers in various plastic culture formats, as well as scalable production of 3D airway organoid. Furthermore, in line with previously published human airway organoid models ^11,12^, we validated submerged-differentiated human nasal epithelial cells as models for PCD, CF, and RSV infection.

## Results

### Notch and BMP inhibition promotes the differentiation of submerged-cultured nasal epithelia

Hypoxic conditions in submerged cultures are associated with suppressed airway epithelial differentiation and enhanced activation of Notch and BMP signaling pathways ^13–15^. To promote differentiation in submerged nasal airway epithelial cultures, we therefore evaluated the effects of the Notch-targeting γ-secretase inhibitor DAPT and the BMP inhibitor DMH1, in combination with a previously established differentiation medium used for ALI cultures ^16,17^.

Submerged differentiation was assessed using cryopreserved human basal cells (BCs) derived from nasal brushings of healthy donors. These cells were cultured as confluent monolayers in conventional 96-well culture plates and differentiated for 21 or 42 days (Fig. 1a). In contrast to bronchial epithelial cells ^13^, Notch inhibition alone was insufficient to induce the differentiation of ciliated cells in nasal cultures. However, co-treatment with DAPT and DMH1 significantly increased the number of β-tubulin IV^+^ ciliated cells in submerged-differentiated human nasal epithelial cells (S-diff HNEC) compared to individual treatments (Fig. 1b, c). The number of MUC5AC^+^ secretory cells remained consistent across conditions, indicating that secretory cell differentiation occurs effectively in submerged cultures under standard ALI differentiation conditions. This correspondence with a previous study, reporting persistent MUC5AC^+^ secretory cell differentiation in ALI-cultured HNEC under hypoxic conditions ^18^. DAPT combined with the BMP inhibitor Noggin also enhanced ciliated cell differentiation, whereas BMP4 co-stimulation inhibited this process (Supplementary Fig. 1a, b).

**Figure 1:**
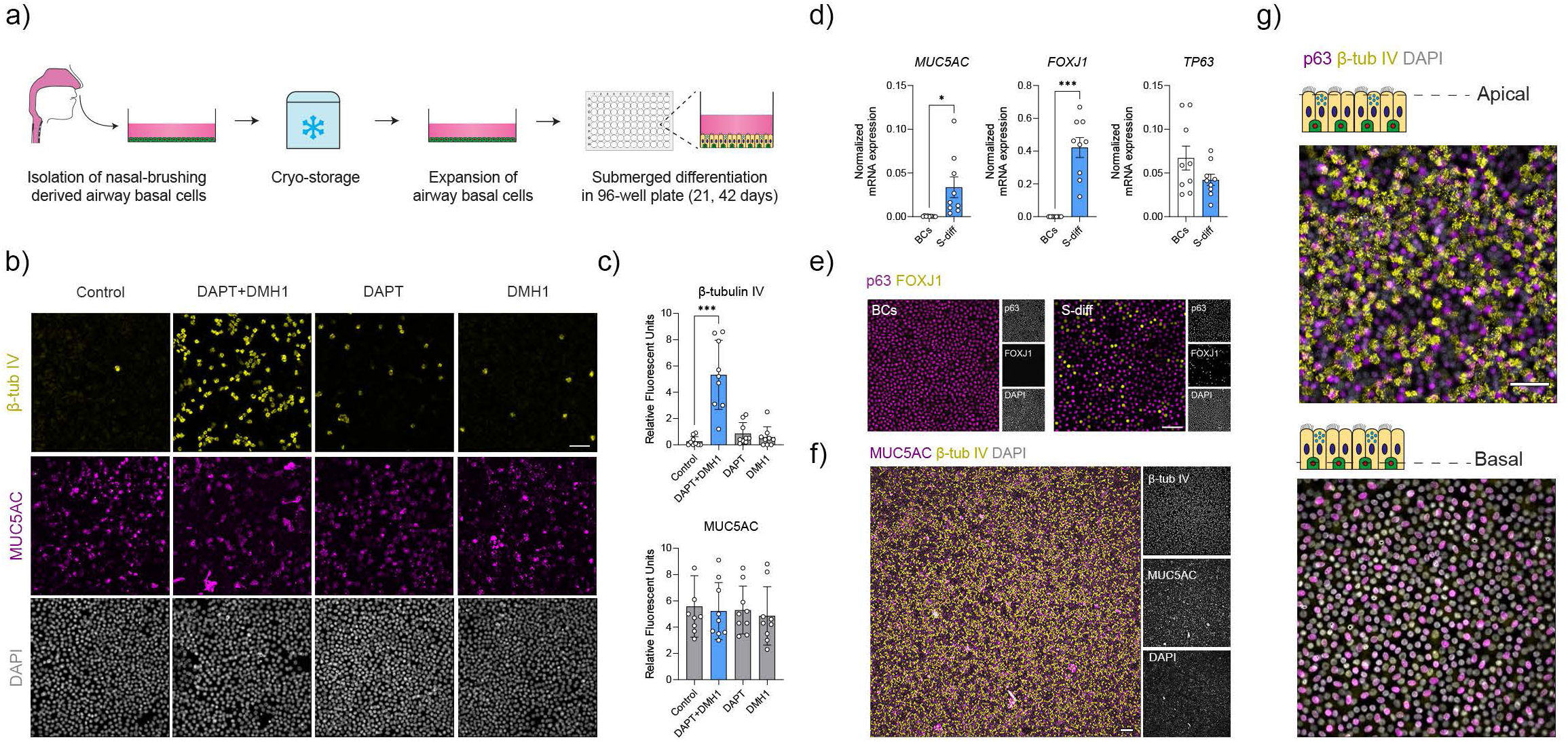
Submerged differentiation of human airway epithelia with DAPT and DMH1. a) Graphic illustration showing the workflow of expanding cryo-stored nasal brushing-derived human upper airway basal cells, followed by experiments investigating differentiation in submerged cultures. b) Representative immunofluorescent images of randomly selected S-diff HNEC after differentiation with DAPT and DMH1 for 21 days. Cells were fixed and stained for the ciliated cell marker β-tubulin IV (β-tub IV; yellow), secretory cell marker MUC5AC (purple), and DAPI (gray). c) Quantification of β-tubulin IV (β-tub IV) and MUC5AC signal (n=3 images for 3 independent donors). d) Quantitative PCR comparing the expression of *MUC5AC*, *FOXJ1,* and *TP63* between BCs and S-diff HNEC of healthy donors (n=2 replicates for 9 independent donors). e) Representative immunofluorescent images of BCs and S-diff HNEC stained for the ciliated cell transcription factor FOXJ1 (yellow), basal cell transcription factor p63 (purple), and DAPI (gray). f) Representative immunofluorescent images of S-diff HNEC differentiated for 42 days stained for β-tubulin IV (β-tub IV; yellow), MUC5AC (purple), and DAPI (gray). g) Representative immunofluorescent images of S-diff HNEC differentiated for 42 days, demonstrating β-tubulin IV (β-tub IV; yellow) staining at the apical side (upper panel), and p63 staining (purple) located more at the basal side of the culture. Data are presented as mean ± SD, and individual data points. Statistical significance was tested using (c,d) a two-tailed paired t-test. *: p<0.05, and ***: p<0.001.

S-diff HNEC displayed significantly higher mRNA expression of *MUC5AC* and the ciliated cell-associated transcription factor *FOXJ1* compared to undifferentiated BCs, while expression of the basal cell transcription factor *TP63* remained unchanged (Fig. 1d). Immunofluorescence staining confirmed these findings, showing that S-diff HNEC, but not BCs, contained both FOXJ1^+^ and p63^+^ nuclei (Fig. 1e). In addition to MUC5AC, secretory epithelial markers SLPI, pIgR, and CC16 (SCGB1A1) were detected in S-diff HNEC (Supplementary Fig. 1c). S-diff HNECs maintained for up to 42 days in culture without passaging exhibited uniform cellular distribution within the epithelial monolayer (Fig. 1f), with ciliated cells localized at the apical side and basal cells at the basal side (Fig. 1g), recapitulating the spatial organization of native airway epithelia. In summary, these results demonstrate that inhibition of Notch and BMP signaling effectively promotes the differentiation in submerged nasal airway epithelial cultures.

### Submerged-differentiated cultures display airway epithelial heterogeneity

To further characterize the cellular heterogeneity of S-diff HNEC, we performed single-cell RNA sequencing (scRNA-seq) using the SORT-seq protocol ^19^, analyzing HNEC differentiated in 6-well plastic culture plates for 21 days (Fig. 2a).

**Figure 2:**
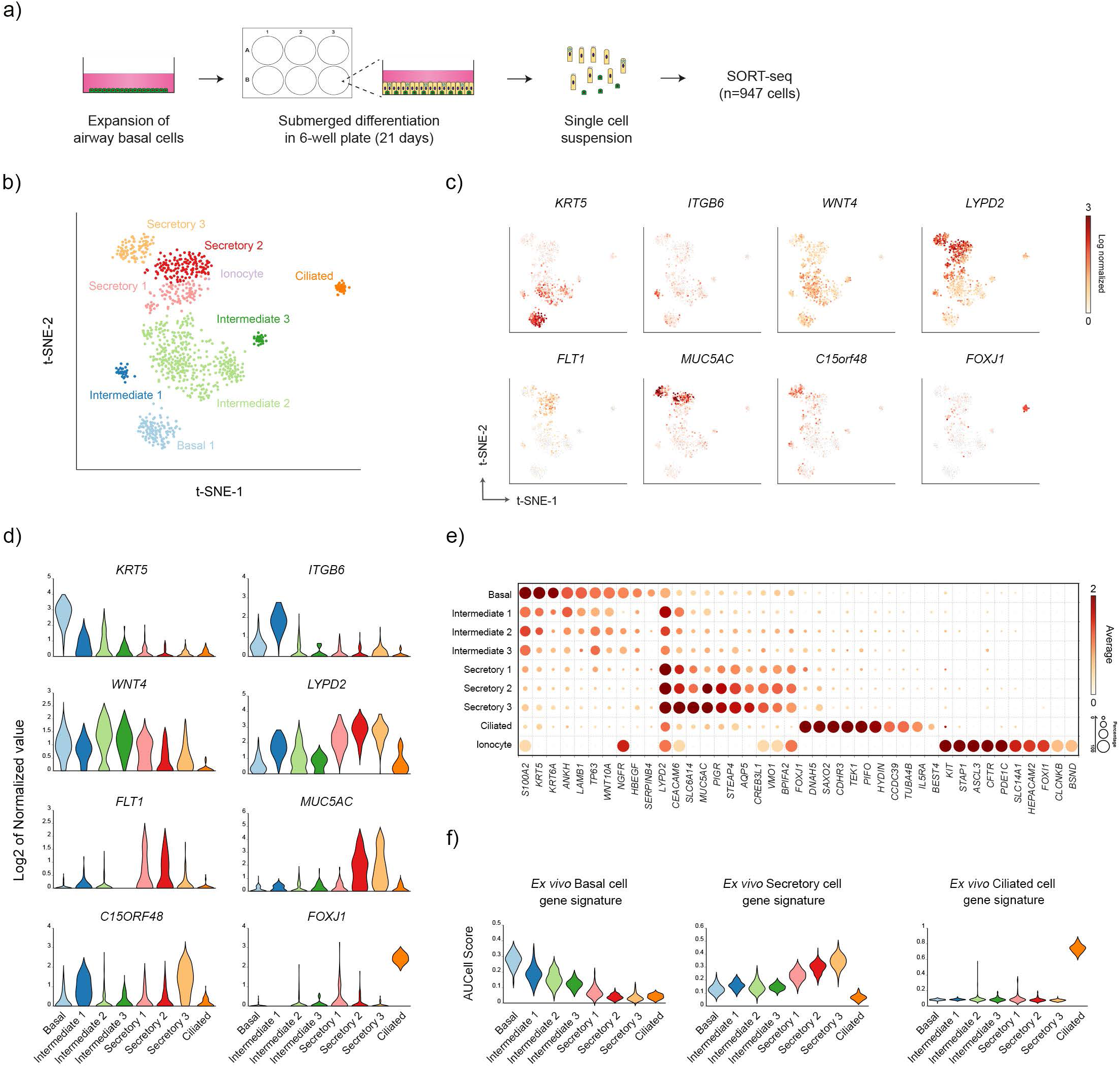
Single cell RNA sequencing analysis of submerged-differentiated airway epithelia. a) Graphic illustration showing the workflow of single cell RNA sequencing experiments with S-diff HNEC. b) t-SNE analysis of expression from scRNA-seq (947 cells in total) of HNEC (n=3 independent donors) which were differentiated in submerged cultures for 21 days. c) Clusters are labelled in t-SNE plots by cell identity based on marker gene expression. d) Violin plots showing the expression of marker genes of identified cell subsets in scRNA-seq data. e) Bubble heatmap showing the expression of selected marker gene expression of identified cell types in S-diff HNEC. f) AUCell scores of *ex vivo* basal, secretory, and ciliated cells gene signatures in S-diff cultures.

Here, we observed the presence of BCs, intermediated, secretory, and ciliated cells (Supplementary table 1), which were further classified into nine distinct clusters representing airway epithelial subsets (Fig. 2b). One cluster of BCs exhibited high expression of *KRT5*, compared to three intermediate subsets that displayed lower expression of basal cell markers (Fig. 2c-e). These intermediate subsets were characterized by elevated *WNT4* expression (Fig. 2c and d), previously implicated in promoting ciliated cell differentiation ^20^. Notably, we identified an intermediate subset marked by high *ITGB6* expression (Fig. 2c and d), resembling basaloid-like cells described in ALI-differentiated cultures ^21^.

Secretory cells, defined by high *LYPD2* expression and other secretory markers, were classified into three distinct subsets (Fig. 2c-e). This included one subset lacking *MUC5AC* but expressing VEGFR receptor 1 (*FLT1*), and two *MUC5AC* expressing subsets (Fig. 2c and d). One *MUC5AC*^+^ subset was characterized by the expression of *C15ORF48*, a marker previously also identified in scRNA-seq analyses of native nasal airway epithelia ^22^. This subset displayed elevated expression of secretory markers, suggesting a more mature secretory cell phenotype compared to other subsets (Fig. 2e). A distinct ciliated cell cluster was also identified, with high expression of *FOXJ1* and other cilia-related genes (Fig. 2c-e). Despite the presence of these major airway epithelial cell types, we detected only a single ionocyte ^23^, likely due to the limited number of cells analyzed (Fig. 2e).

To further validate the identity of major cell clusters in S-diff cultures, we cross-referenced the expression of basal, secretory, and ciliated cell gene signatures derived from *ex vivo* nasal airway epithelium from the integrated human lung atlas ^24^ (Fig 2f). Calculation of AUCell scores confirmed enrichment of corresponding subsets in S-diff HNECs. Collectively, these results confirm the presence of major airway epithelial cell types and demonstrate the presence of cellular heterogeneity of S-diff HNEC cultures.

### Comparable transcriptomes between S-diff and ALI-differentiated HNEC

To assess transcriptomic similarities between S-diff HNEC, undifferentiated BCs, and ALI-differentiated cultures, we performed bulk RNA sequencing (Fig. 3a). In alignment with submerged cultures, the differentiation medium of ALI cultures was supplemented with DAPT and DMH1, which resulted in a higher proportion of ciliated cells compared to S-diff HNEC (Supplementary Fig. 2a).

**Figure 3:**
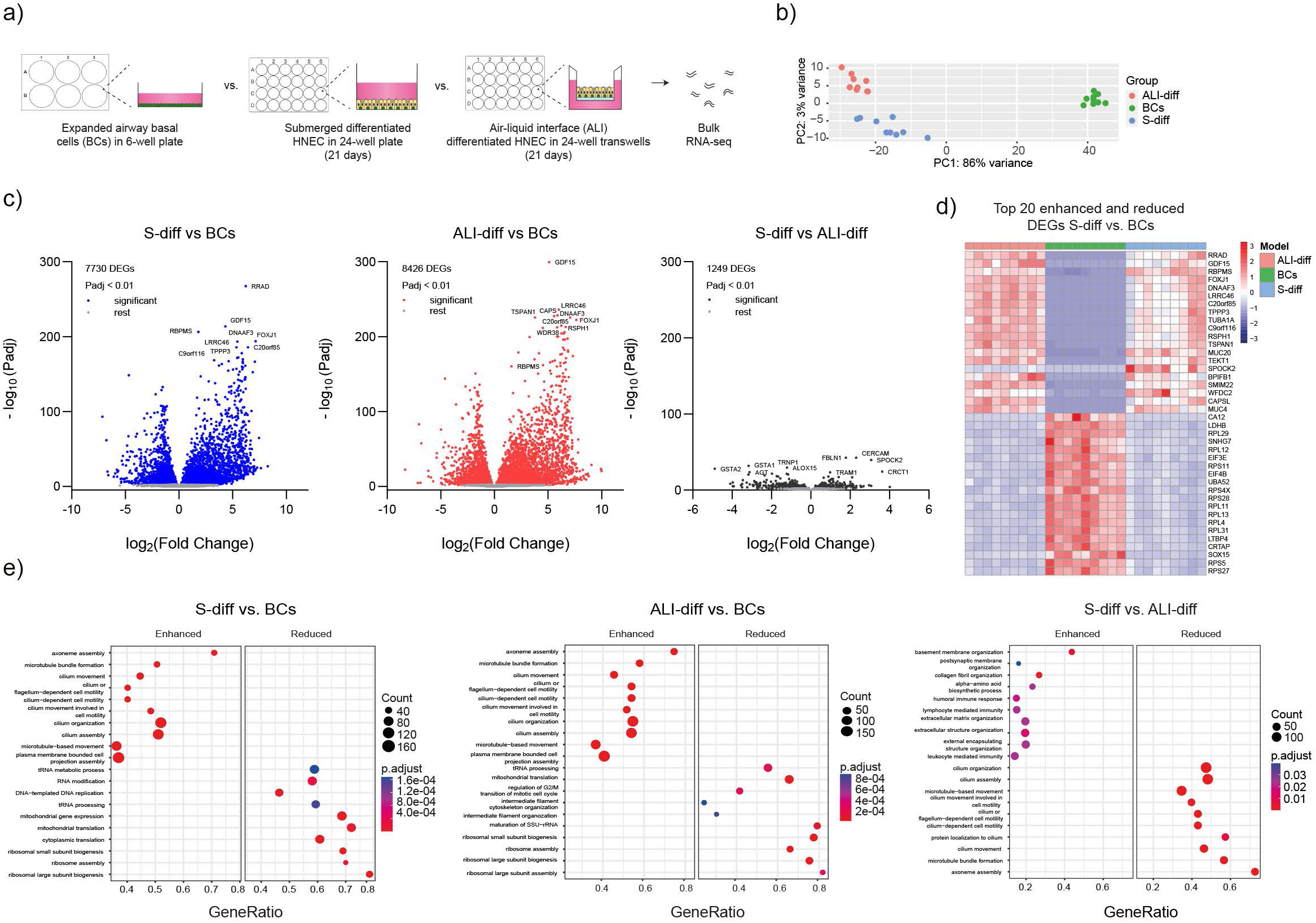
Comparative analysis of S-diff HNEC with BCs and ALI-differentiated HNEC. a) Graphic illustration showing the use of expanded airway basal cells, S-diff and ALI-differentiated HNEC for RNA-sequencing (RNA-seq) (n=9 independent donors). b) PCA plot from bulk RNA-seq data, comparing S-diff HNEC with basal cells (BCs) and ALI-differentiated cultures. c) Volcano plots displaying the log_2_ fold change and the log_10_ differentially expressed genes (DEGs) in S-diff HNEC compared to BCs (left panel), ALI-differentiated cultures compared to BCs (middle panel), and S-diff HNEC compared to ALI-differentiated cultures (right panel). For all three comparisons, the top 10 DEGs are mentioned by name. d) Heatmap showing marker gene expression of top 20 enhanced and reduced DEGs of S-diff HNEC vs BCs. e) GO-term analysis of the top 10 activated and suppressed biological processes in S-diff vs. BCs (left panel), ALI-differentiated vs. BCs (middle panel), and S-diff vs ALI-differentiated HNEC (right panel).

Principal Component Analysis (PCA) of bulk RNA-seq data revealed that S-diff HNEC exhibit a transcriptome more similar to ALI-differentiated HNEC than to BCs, with 3% and 86% variance, respectively (Fig. 3b). Further analysis identified 7,730 differentially expressed genes (DEGs) between S-diff HNECs and BCs (P < 0.01), while 1,249 DEGs were observed between S-diff and ALI cultures (Fig. 3c, Supplementary Fig. 2b, Supplementary Table S2). Notably, the DEGs observed in S-diff HNEC vs. BCs overlapped substantially with those identified between ALI-differentiated cultures and BCs (Fig. 3c, d). Analysis of airway epithelial cell subset-related genes confirmed significantly lower expression of BC-related markers and higher expression of secretory and ciliated-cell markers in both S-diff and ALI-differentiated HNEC compared to BCs (Supplementary Fig. 2c). However, S-diff HNEC showed significantly lower expression of ciliated-cell markers compared to ALI cultures. Gene ontology (GO) term analysis demonstrated that both S-diff and ALI cultures shared enriched gene sets related to cilia structure and function, while reduced gene sets included pathways associated with transcription and translation. Differences between S-diff and ALI cultures were observed in pathways related to ciliary function, consistent with the higher abundance of ciliated cells in ALI cultures (Fig. 3e). In contrast, S-diff cultures showed an enrichment of gene sets related to the basal membrane, extracellular matrix, and immune responses compared to ALI cultures (Fig. 3e, Supplementary Fig. 2d).

Due to the relatively low number of ciliated cells, we extended the differentiation period to evaluate its effects (Supplementary Fig. 3a). *FOXJ1* mRNA expression and IF staining of β-tubulin IV was highest at day 42 (Supplementary Fig. 3b-d). This was paired with a decline in MUC5AC expression, suggesting trans-differentiation of secretroy cells into ciliated cell as late-stage mechanism. Moreover, the expression of the ionocyte-related transcription factor *FOXI1*, as well as *CFTR*, was highest at day 42 (Supplementary Fig. 3b).

Altogether, bulk RNA-seq data suggest that S-diff HNEC are more comparable to ALI-differentiated HNEC when compared to undifferentiated BCs. Extending the differentiation period in S-diff cultures increases the number of ciliated cells, further aligning their cellular composition with ALI-differentiated cultures.

### Cilia dynamics of submerged-differentiated cultures of healthy controls and PCD subjects

As an initial demonstration of the suitability of S-diff HNEC for studying respiratory diseases, we investigated its potential in the context of PCD. This heterogeneous monogenic disorder is characterized by airway epithelial ciliopathy, with mutations reported in approximately 50 different PCD-related genes ^8^. These mutations can lead to cilia immotility or abnormal wave patterns in motile cilia, reduced assembly of cilia structures, or attenuated ciliated cell differentiation.

First, we compared cilia motility in S-diff cultures (Video S1) to ALI-cultures (Video S2) of healthy controls (HC) by measuring the ciliary beat frequency (CBF), which did not differ significantly (Fig. 4a,b). Monitoring cilia motility during the differentiation of submerged cultures demonstrated a stable CBF between 21 and 42 days (Supplementary Fig. 4a). Next, we measured the CBF in S-diff HNEC from PCD subjects with distinct genotypes (Fig. 4c and d). In line with previous studies ^25,26^, we did not detect ciliary movement in PCD subjects with mutations in *DNAH5* (Video S3 and S4). A significantly lower CBF was observed in S-diff HNEC with *HYDIN* mutations (Video S5), when compared to HCs. In S-diff HNEC with mutations in *CCDC40* ^27^, a fraction of the ciliated cells displayed motile cilia (Video S6) with a lower CBF compared to HCs. In line with impaired generation of cilia ^28^, S-diff HNEC of a PCD subject with *CCNO* mutations displayed reduced ciliary abundance, whereas FOXJ1 protein was abundantly present (Fig. 4e).

**Figure 4:**
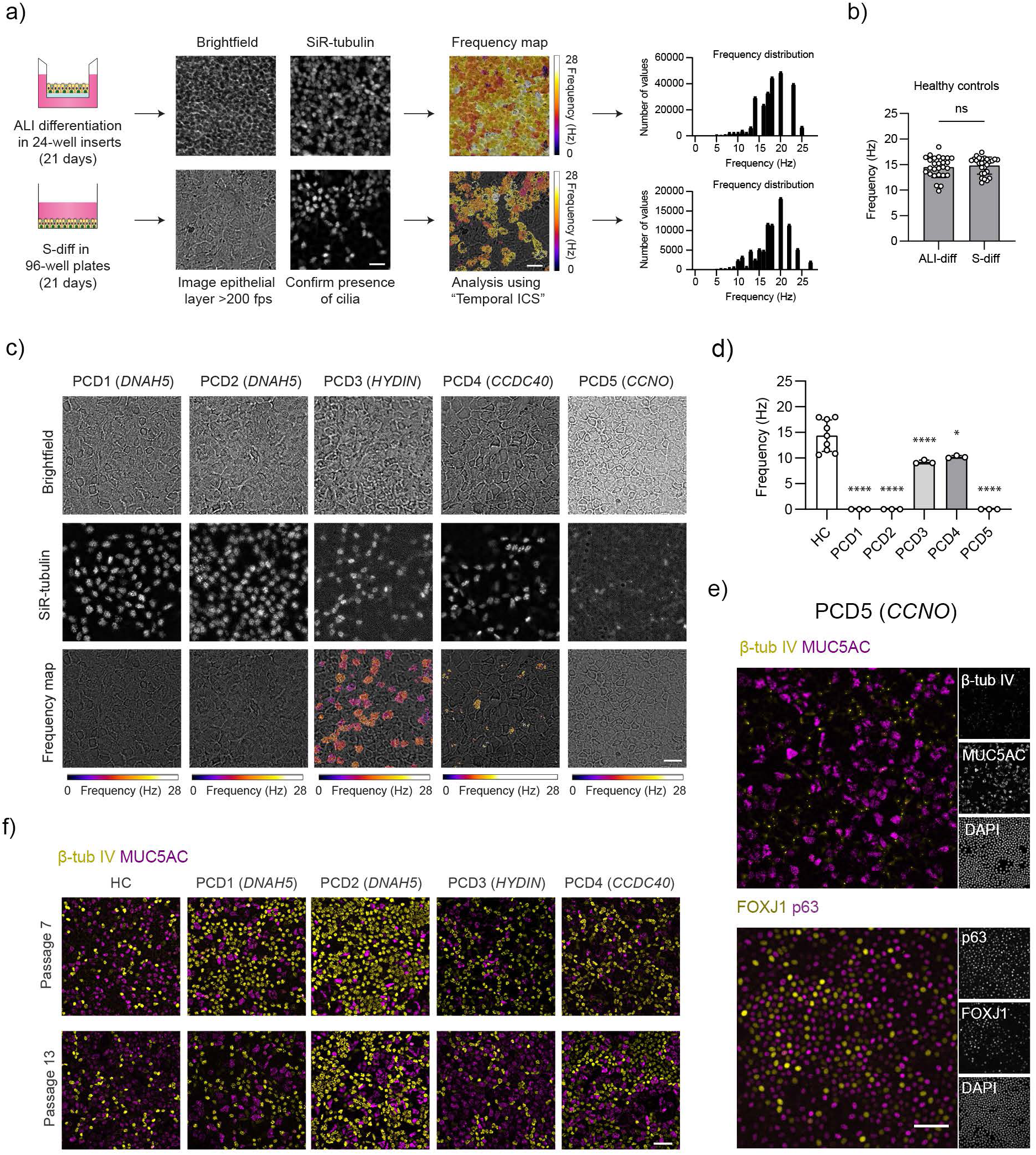
Ciliary beat frequency (CBF) in S-diff HNEC of healthy controls and PCD subjects. a) Graphic illustration showing the workflow of ciliary beat frequency (CBF) measurements. Representative brightfield and immunofluorescent video frames of ALI- and S-diff HNEC incubated with the live dye SiR-tubulin with corresponding frequency map generated with “Temporal ICS” analysis. Video frames were extracted from Video S1 and S2. Scale bar: 50 µm. Frequency maps show CBF in hertz (Hz). Color bar indicates CBF from 0-28 hertz (Hz). b) Average CBF in Hz of submerged and ALI-differentiated cultures (n=3 video’s for 9 independent healthy donors). Each point represents the mean frequency for one video. c) Representative bright-field and immunofluorescent video frames of submerged-differentiated PCD cultures (PCD1-5) incubated with the live dye SiR-tubulin with corresponding frequency maps and measured ciliary beat frequency (CBF). Frequency is presented in hertz (Hz). Color bar indicates CBF from 0-28 hertz (Hz). d) CBF in Hz of submerged-differentiated healthy (n=9) and PCD (n=5, PCD1-5) donors. Each point represents the mean frequency for one video. e) Representative immunofluorescent image of S-diff HNEC of a PCD donor with mutations in CCNO (PCD5) stained for β-tubulin IV (β-tub IV; yellow), MUC5AC (purple) (Top panel), p63 (purple), and FOXJ1 (yellow) (bottom panel), and DAPI (gray) f) Representative immunofluorescent images of p7 and p13 S-diff HNEC of one healthy donor and PCD subjects PCD1-4. Cells were fixed and stained for the secretory cell marker MUC5AC (purple) and ciliated cell marker β-tubulin IV (yellow). Data are presented as mean ± SD, and individual data point. Statistical significance was tested using a (b) two tailed paired t-test, (d) Tukey’s multiple comparison test. ns = non-significant, *: p<0.05, ****: p<0.0001.

To validate prolonged use of S-diff HNEC in personalized screening assays, we examined BCs from HC and PCD donors, expanded up to passage 6 and 12 in feeder-free expansion conditions (Supplementary Fig. 4b). We observed consistent population doublings and the absence of morphological differences until passage 12 in all examined donors (Supplementary Fig. 4c and d). MUC5AC and β-tubulin IV staining of S-diff HNEC were similar between passage 7 and 13 in HC donors, however significant differences were observed in PCD donors (Fig. 4f, Supplementary Fig. 4e). Despite this variability, the average CBF of S-diff HNEC at passage 7 and 13 were comparable for both HC and PCD donors (Supplementary Fig. 4f) and corresponded with observations at passage 4 (Fig. 4d). Collectively, we demonstrated comparable CBF measurement in ALI- and S-diff HNEC and genotype-dependent ciliopathy in cultures of individuals with PCD, which could be consistently measured in S-diff HNEC after long-term expansion of BCs.

### CFTR modulator responses in CF nasal airway organoids generated from S-diff monolayers

Next, we investigated the application of S-diff HNEC as CF respiratory disease model. CF is caused by autosomal recessive inherited mutations in the cystic fibrosis transmembrane conductance regulator (CFTR) gene, in which impaired CFTR protein function in airway epithelia leads to reduced anion and fluid secretion, resulting in the dehydration of secreted mucins that cannot be removed by motile cilia ^29^. CFTR modulating drugs, i.e. correctors and potentiators, can restore CFTR function ^30^. However, drug efficacy depends on CFTR genotype, which varies between individuals with CF.

We previously described a method for scalable generating of uniform-differentiated airway organoids from ALI-differentiated monolayer fragments ^16,17^. These organoids can be used to examine CFTR modulator efficacy in forskolin-induced organoid swelling (FIS) assays. Here, we aimed to explore whether S-diff HNEC cultured in more scalable culture dishes could also be converted into organoids and used in FIS assays to study CFTR modulator responses (Fig. 5a). HNEC were differentiated in thermosensitive dishes, which enabled epithelial dissociation by placing the dish on a cold surface. When embedded in an extracellular matrix, epithelial fragments of S-diff cultures self-organized into cystic organoids within 48 hours (Fig. 5b). IF staining of β-tubulin IV, MUC5AC, and p63 confirmed the presence of different airway epithelial subsets (Fig. 5c).

**Figure 5:**
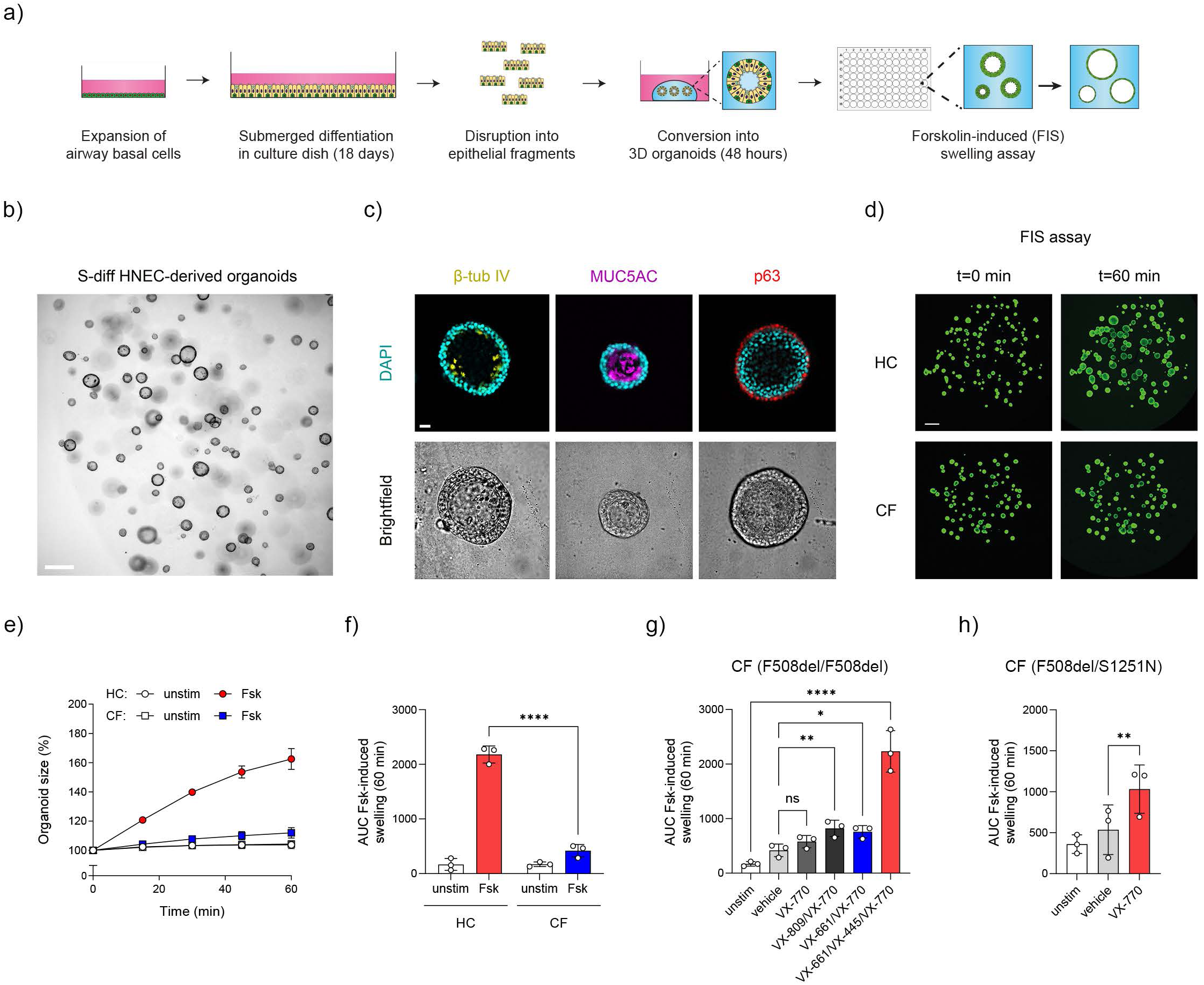
Conversion of S-diff HNEC into 3D organoids and CFTR function measurements. a) Graphic illustration showing the workflow of generating airway organoids from epithelial fragments of S-diff HNEC, and application in forskolin-induced swelling assays. b) Representative brightfield images of S-diff monolayer derived cystic airway organoids formed after 48 hours. c) Immunofluorescent (top) and brightfield (bottom) images of S-diff HNEC-derived organoids of β-tubulin IV (yellow, left top), MUC5AC (purple, middle top), p63 (red, right top), and DAPI (cyan). d) Representative images of forskolin-induced swelling determined with calcein green AM esters–stained healthy control (HC) and cystic fibrosis (CF; F508del/F508del) organoids, with images taken before stimulation with forskolin (t=0, left panels) and 60 minutes after stimulation (t=60 min, right panels). e,f) Comparison of FIS between HC and CF airway organoids derived from S-diff HNEC (both n=3 independent donors) that were unstimulated or stimulated with forskolin (Fsk). g) CF F508del homozygous organoids derived from S-diff HNEC (n=3 independent donors) were pre-treated with vehicle or CFTR correctors, VX-809, or VX-661/VX-445 for 48 hours. Afterwards FIS was determined following acute stimulation with Fsk, VX-770, or vehicle. h) CF F508del/S1251N airway organoids generated from S-diff HNEC (n=3 independent donors) were used in FIS assays, together with acute stimulation with the CFTR potentiator VX-770. FIS results are depicted as (e) the percentage change in surface area relative to t = 0 (normalized area) measured at 15-min time intervals for 60 minutes, or (f) as area-under-the-curve (AUC) plots (t = 60 min). Data are presented as mean ± SD, and individual data point. Statistical significance was tested using a (f) two tailed paired t-test, (g,h) Tukey’s multiple comparison test. ns = non-significant, *: p<0.05, **: p<0.01. ****: p<0.0001.

FIS assays were conducted in organoids stimulated with inflammatory mediators to increase CFTR function, as previously described ^16^. In these conditions we observed a significantly lower swelling in CF organoids compared to HCs, consistent with impaired CFTR function (Fig. 5d-f). Moderate-to-high increases in swelling were observed in CF F508del/F508del organoids with the CFTR modulator combinations VX-809/VX-770, VX-661/VX-770, and VX-661/VX-445/VX-770 (Fig. 5g). Furthermore, the CFTR potentiator VX-770 increased swelling of organoids from individuals with CF carrying a single S1251N gating mutation (Fig. 5h). Altogether, these results support that S-diff monolayers in scalable dishes can effectively be converted into differentiated organoids and used as CF personalized disease model.

### RSV infections in submerged-differentiated airway epithelial cells

Next, we used S-diff HNEC to study infections with RSV (Fig. 6a), which imposes a significant health burden on vulnerable populations such as infants in low-to-middle income countries and individuals with chronic respiratory diseases ^9^.

**Figure 6:**
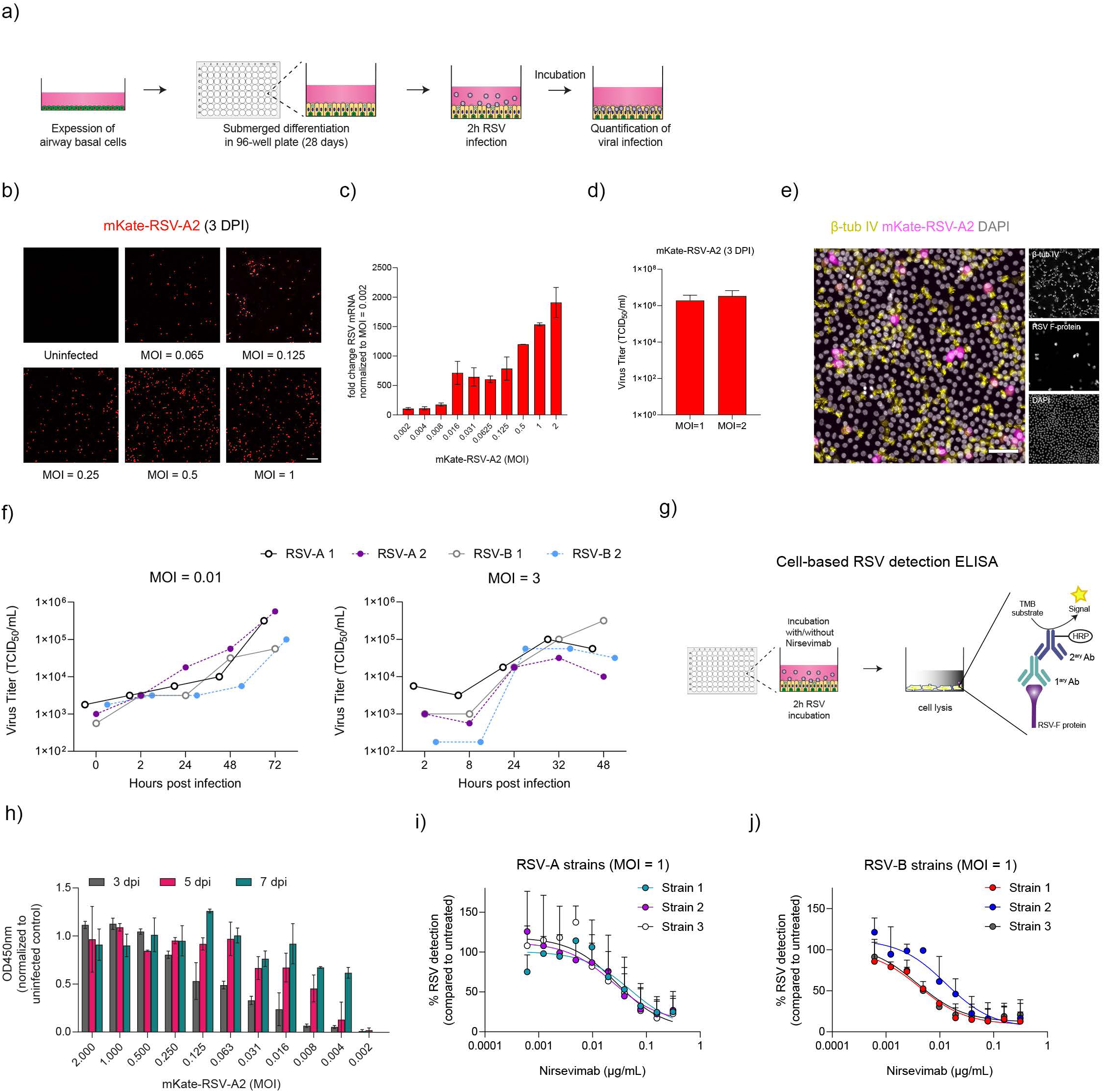
Modeling respiratory syncytial virus infections in S-diff HNEC. a) Graphic illustration summarizing experimental workflow of RSV infections in 96-well plate cultures S-diff HNEC. b) Live imaging of RSV-A2 mKate (red) infecting S-diff HNEC underlines an MOI-dependent infection at 3 DPI. c) RSV mRNA levels in mKate-RSV-A2-infected S-diff HNEC at 3 DPI. Data is shown as fold change, normalized to the lowest MOI (0.002) and housekeeping genes. d) Infectious viral titers reported as TCID50 per mL for RSV-A2 mKate using a TCID50 assay. S-diff HNEC were infected with RSV-A2 mKate at MOI=2 and MOI=1 and subsequently viral supernatants were collected and tittered with a TCID50 assay at 3 DPI. e) Representative immunofluorescent images of S-diff HNEC infected with mKate-RSV-A2 (magneta) of β-tubulin IV (yellow), and DAPI (cyan). f) S-diff HNEC were infected with clinical RSV-A and -B strains at an MOI = 0.01 (left) or MOI = 3 (right). Virus growth kinetics was determined by titrating supernatant and cell-bound RSV at different time intervals following infection. g) Graphic illustration describing the concept of the cell-based ELISA used with S-diff HNEC to quantify RSV infections. h) S-diff HNEC were infected with RSV-A2 mKate at an MOI range and RSV infectivity was measured and depicted as OD450 nm corrected to uninfected control after 3, 5 and 7 DPI. i,j) Assessment of the neutralization activity of Nirsevimab against clinical (i) RSV-A and (j) RSV-B strains. RSV infections in S-diff HNEC were conducted with a MOI=1 at 3 DPI. RSV infections were quantified by ELISA and depicted as the percentage RSV detection compared to uninfected controls. Experiment were conducted 2-3 times in S-diff HNEC of one healthy donor. Data are presented as mean ± SD.

First, we infected S-diff cultures with an mKate-RSV-A2 strain, observing an MOI-dependent increase in mKate-fluorescence at 3 days post-infection (DPI) (Fig. 6b). qPCR analysis confirmed RSV mRNA expression in cell lysates (Fig. 6c), and titration of clarified cell pellets on HEp-2 cells demonstrated the production of infectious virions (Fig. 6d). mKate-RSV-A2 fluorescence co-localized in β-tubulin IV^+^ cells (Fig. 6e), consistent with tropism for ciliated cells. Comparative studies demonstrated higher RSV infection in the ALI-model (Supplementary Fig. 5a), in line with a higher abundance of ciliated cells. Further infection studies were conducted with RSV-A and -B clinical-isolate strains demonstrating a time-dependent increase in the production of infectious virions in both S-diff HNEC and HEp-2 cells (Fig. 6f, Supplementary Fig. 5b). Furthermore, under a high viral infection condition (MOI=3), a plateau in production of infectious virions was reached, indicating saturation of infectivity.

Matching the scalable character of the S-diff culture model, we optimized a cell-based ELISA, to simplify and increase the throughput of RSV infection measurements (Fig. 6g). With the ELISA, we observed MOI-dependent RSV detection after infection with mKate-RSV-A2 in S-diff HNEC at 3 DPI (Fig. 6h). Furthermore, at MOIs ranging from 0.004-0.125 a time dependent increase in RSV detection was observed, while at higher MOIs RSV detection was already saturated at 3 DPI. Validation of the ELISA demonstrated adequate assay performance (coefficient of variation values: 12.7% and 9.4%, respectively), and the ability to discriminate noise and signal (Z’-factor of 0.48) (Supplementary Fig. 5c). Moreover, we observed a positive correlation (rs2=0.73, p<0.005) with the qPCR-based quantification of RSV F-mRNA in S-diff HNEC (Supplementary Fig. 5d). With the ELISA, we observed considerably lower RSV infections in S-diff HNEC than in HEp-2 and VERO E6 cells, confirming previous studies describing higher levels of infection in transformed cell lines ^31^ (Supplementary Fig. 5e).

Lastly, we confirm the application of the cell-based ELISA with S-diff HNEC in viral neutralization experiments with the RSV-neutralizing monoclonal antibody Nirsevimab ^32^. Here, we observed a dose dependent decrease in the detection of RSV in infection experiments with mKate-RSV-A2, RSV-A, and -B clinical-isolate strains (Fig. 6i,j, and Supplementary Fig. 5f). Consistent with previous studies in other model systems ^32^, we furthermore observed a near-complete inhibition at saturating concentration and an IC50 of 2.90 μg/mL. In summary, we demonstrate the feasibility of using S-diff HNEC in RSV infections and viral neutralization experiment together with a cell-based RSV detection ELISA.

## Discussion

In this study, we characterized small-molecule directed differentiation of submerged-cultured human airway epithelia and explored applications as a respiratory disease model. Previous studies supported the feasibility of submerged differentiation of human bronchial and murine airway epithelial cells ^13,33^. Our approach is distinct in its specific use of nasal epithelial cells, obtained through nasal brushings, to enable personalized disease modeling of monogenic respiratory disorders and upper airway viral infections.

A key advantage of S-diff HNEC is the flexibility and cost-efficiency of using conventional culture plastics. In addition, S-diff monolayers from large plastic surfaces can be used to generate 3D organoids, reducing the use of basement membrane extract normally required for organoid expansion and improving the efficiency of culturing uniformly differentiated airway organoids at scale. Importantly, S-diff HNEC more closely resemble the native human airway epithelium and ALI-differentiated cultures compared submerged cultures of undifferentiated primary airway epithelia and transformed cell lines. Therefore, we propose a novel workflow where 2D S-diff cultures or 3D organoids generated from these monolayers serve as practical alternative to traditional cell culture models, particularly for large-scale and miniaturized applications. Findings from the S-diff model can be further validated in more complex model systems such as ALI-differentiated airway epithelial cells.

We validated the application of S-diff HNEC for studying the monogenic respiratory diseases PCD and CF, demonstrating their ability to recapitulate key disease characteristics. S-diff HNEC hold further potential for diagnostic purposes and personalized screening of novel therapies for both PCD and CF, such as readthrough therapies targeting stop codon mutations, as well as gene- and mRNA-based therapies using lipid nanoparticles or other delivery methods ^34,35^. In our validation studies on RSV infection, we optimized a scalable cell-based ELISA for higher-throughput quantification, compatible with conventional 96-well plates. This method, which is independent of exogenous fluorescent protein expression in lab strains, allows higher-throughput characterization of clinical isolate strains. We propose that S-diff HNEC could be further applied to examine RSV evolution, identify viral escape variants to clinically approved therapies, and to characterize novel antiviral therapies ^9^. Additionally, this model could be used for personalized viral infection studies, such as in individuals with immune deficiencies due to genetic mutations, as demonstrated in prior studies with ALI-differentiated nasal epithelia from RSV-susceptible individuals with CD14 deficiency. ^10^.

While our validation studies mainly focused on ciliated and secretory cells, further characterization of basal cells and rare cell types in S-diff cultures remain needed. Moreover, small molecules targeting additional signaling pathways may further enhance the differentiation in submerged cultures. This includes optimizing ciliated cell differentiation to levels similar to the ALI-model or enriching rare cell types, for instance with small molecules targeting the Sonic Hedgehog pathway to increase ionocytes ^36^. Other potential application of S-diff HNEC include studying biological pathway involved in ciliated cell differentiation or motile cilia function, employing CRISPR gene editing screening assays, or developing co-culture models with immune cells to test anti-inflammatory therapies for chronic inflammatory airway diseases using patient-derived cells.

In conclusion, we introduce submerged-differentiated human upper airway nasal epithelia as versatile model for respiratory disease research, combining the simplicity of traditional cell cultures with the cellular complexity of the human airway epithelium. Similar to human airway organoids ^11,12^, we provide proof-of-concept of using S-diff cultures as model for PCD, CF and RSV infection. Beyond these application, the S-diff model holds promise for scaling up *in vitro* studies in cell biology, personalized medicine, and drug discovery for other respiratory diseases. Additionally, the small-molecule-directed differentiation methodology used in submerged culture conditions could be adapted to other epithelial tissues, indicating broader application of this simple and flexible culture method in human cellular disease modeling.

## Declaration of generative AI and AI-assisted technologies in the writing process

During the preparation of this work the authors used ChatGPT 4.0 in order to improve language and readability. After using this tool, the authors reviewed and edited the content as needed and take full responsibility for the content of the publication.

## Acknowledgments

This study was supported by a Health∼Holland Top Consortium Knowledge and Innovation (TKI) grants (LSHM18062), grants of the Dutch Cystic Fibrosis Foundation (NCFS, HIT-CF grant), the Netherlands Organization for Scientific Research (NWO) Gravitation programme IMAGINE! (project number 24.005.009), and funding from AstraZeneca.

## Author Contributions

Conception and/or design: HD, GI, SS, LB, JB, GA; Acquisition, analysis, or interpretation of data: HD, GI, SS, MI, EK, JT, LR, LH, SS, IW, LA, LB, KP, SB, RL EH, CE, LK, JB, GA; Drafting the work or revising it critically for important intellectual content: HD, GI, SS, LB, JB, GA; Final approval of the version submitted for publication: HD, GI, SS, MI, EK, JT, LR, LH, SS, IW, LA, LB, KP, SB, RL EH, CE, LK, JB, GA.

## Methods

### Human materials and informed consent

Nasal brushings from healthy volunteers (n=10 independent donors), PCD subjects (n=5 independent donors), and CF subjects (n=6 independent donors) were collected as previously described ^17^. All donors gave informed consent and this study was approved by the Institutional Medical Research Ethics Committee of the University Medical Center Utrecht (Toetsingscommissie Biobank Utrecht, the Netherlands) protocol ID: 16-586, and 21-044.

### Isolation and expansion of airway epithelial basal cells from nasal brushings

Nasal brushing-derived airway epithelial basal cells were isolated essentially as previously described ^17^. In brief, dissociated single cells were seeded in a collagen IV (Cat#C7521, Sigma)-precoated cell culture plate (6-well; Cat#657160, Greiner Bio-One) and cultured in isolation medium consisting of 50% (v/v) BEpiCM-b (Cat#3211, Sciencell) and 44% (v/v) advanced DMEM/F-12 (Ad-DF; Cat#12634-028, Thermo Fisher Scientific) supplemented with 2% (v/v) B-27 Supplement, serum free (Cat#17504001, Thermo Fisher Scientific), 10 mM HEPES (Cat#15630080, Thermo Fisher Scientific), 1% (v/v) GlutaMAX supplement (Cat#35050-061, Thermo Fisher Scientific), 1% (v/v) penicillin/streptomycin (Cat#15070-063, Thermo Fisher Scientific), 0,5 µg/mL Hydrocortisone (Cat#H0888, Sigma-Aldrich), 1,25 mM N-Acetyl-L-cysteine (Cat#A9165, Sigma-Aldrich), 100 µg/mL Primocin (Cat#ant-pm-2, InvivoGen), 1 µM ALK5 inhibitor A83-01 (Cat#2939/10, Tocris), 0,5 µg/mL (±)-Epinephrine hydrochloride (Cat#E4642, Sigma-Aldrich), 5 µM Y-27632 (Cat#S1049, Selleck Chemicals), 2% RSPO3-Fc Fusion Protein conditioned medium (Cat#R001 - 100 mL, U-Protein Express), 50 nM Human Heregulin-beta 1 (Cat#100-03, PeproTech), 100 ng/mL Recombinant human Fibroblast growth factor 10 (FGF10; Cat#100-26, PeproTech), 25 ng/mL Recombinant human Hepatocyte growth factor (HGF; Cat#100-39H, PeproTech). To prevent microbial infections, the following antibiotics were added during the first week of isolation: 250 µg/mL Amphotericin B (Cat#15290018, Thermo Fisher Scientific), 50 µg/mL Gentamicin (Cat#G1397, Sigma-Aldrich) and 50 µg/mL Vancomycin (Cat#SBR00001, Sigma-Aldrich). After the first week, 5 µM NOTCH inhibitor DAPT (Cat#15467109, Thermo Fisher Scientific) and Rapamycin (Cat#553210-1MG, Sigma) were added to the medium. Cells were cultured at 37°C with 5% CO_2_ and medium was refreshed three times a week until 80-90% confluency was reached. Cells were passaged using TrypLE express enzyme (Cat#12605010, Thermo Fisher Scientific). Passage 1 cells were frozen in CryoStor CS10 (Cat#07930, STEMCELL Technologies) supplemented with 5 µM Y-27632 to create a master cell bank and passage 2 cells were frozen to create a work cell bank. Population doublings (PD) were calculated as PD=3.32×(log(cells harvested/cells seeded)).

### Differentiation of HNEC in submerged and ALI-cultures

Basal cells (BCs; passage 3-12) were cultured on conventional culture plates (CELLSTAR, Greiner Bio-One) which were pre-coated with 30 µg/ml PureCol (Cat#5005, Advanced BioMatrix). For differentiation experiments in submerged cultures, cells were seeded in a density of 0.2 – 0.3 × 10^6^ cells per cm^2^ and cultured in 280 µl per cm^2^ expansion medium until reaching 100% confluency after approximately 5-7 days. Afterwards, culture medium was switched to a differentiation medium consisting of 98.5% (v/v) Ad-DF with 100 nM 3,3′,5-Triiodo-L-thyronine sodium salt (Cat#T6397, Sigma-Aldrich), 0.5 µg/ml hydrocortisone, 0.5 µg/ml (±)-Epinephrine hydrochloride, 50 nM A83-01, 100 nM retinoic acid agonist TTNPB (Cat#16144-1, Cayman), 0.5 ng/mL recombinant human EGF (Cat#AF-100-15, PeproTech) and 1% (v/v) penicillin/streptomycin, which was supplemented with 5 µM DAPT and 5 µM BMP inhibitor DMH-1 (Cat#S7146, Selleck Chemicals). Medium was refreshed three times per week. Cultures were washed with 100 µl PBS for 5 minutes once a week. ALI-differentiation on transwells (Cat#3470, Corning) was conducted as previously described ^17^, with minor changes. In short, 0.2 million BCs were seeded on 24-well transwell inserts and cultured in expansion medium until confluency was reached. Then, medium was switched to differentiation medium supplemented with 500 nM A83-01. Next, apical fluid was removed to create an air-liquid interface. After 3-5 days medium was switched to the final differentiation medium, which consisted of differentiation medium with 5 µM DAPT and 5 µM DMH1.

### Immunofluorescent microscopy

2D differentiated cultures on transwells and on 96-well plastic tissue culture plates were fixed in 4% paraformaldehyde for 15 min, permeabilized in 0.3% (vol/vol) Triton-X-100 (Cat#T8787, Sigma-Aldrich) in PBS for 30 min and treated with blocking buffer, consisting of 1% (wt/vol) BSA, and 0.3% (vol/vol) Triton-X in PBS for 60 min. Primary antibodies (1:500 in blocking buffer; Supplementary Table S3) were incubated for two hours. Afterwards, cells were washed three times with PBS and incubated with secondary antibodies (1:500 in blocking buffer; Supplementary Table S3) and DAPI (1:1000, D9542, Sigma) for 30 minutes in the dark. After three washings with PBS, cultures differentiated on plastic were stored in 150 µl PBS at 4°C and cultures differentiated on transwells were cut from the plastic insert and mounted with ProLong Gold antifade reagent without DAPI (#P36934, Thermo Fisher Scientific) on slides. 3D airway organoids were stained as previously described ^16^. Images were acquired with a Leica THUNDER imager using 5, 10, 20, and 40× dry objectives, and processed using Leica software. Stained surface signal of β-tubulin IV and MUC5AC was quantified using GNU Image Manipulation Program (GIMP; https://www.gimp.org/) by using the color threshold and mask function.

### RNA isolation, quantitative real-time PCR and bulk RNA-sequencing

Total RNA was extracted according to the manufacturer’s protocol (RNeasy kit, QIAGEN). RNA yield was measured with a Nanodrop spectrophotometer. cDNA was generated using the iScript cDNA synthesis kit (Bio-rad) according to the manufacturer’s protocol. Quantitative real-time (qPCR) was performed using iQ SYBR Green Supermix (Cat#1708886, Bio-rad) and a CFX96 real-time detection machine (Bio-rad), and primers described in Supplementary Table S4. Gene expression was calculated using the comparative 2-ΔΔCT method and normalized against the housekeeping genes *ATP5B*, *GAPDH*, and *YWHAZ*. For bulk RNA sequencing, extraction and library preparation followed an adapted version of the CELseq2 protocol ^37^. Sequencing was performed with the Illumina NextSeq (Sequencing depth: STANDARD (10M reads/sample)). Sequencing results were mapped to the human genome (hg38) using R software. Bulk RNA-seq count normalization and differential gene expression were analyzed using the DESeq2 package ^38^. Significantly differentially expressed genes of different sample groups were selected using a log2 fold change ((Padj<0.01 and |log2 Fold change|>1) and adjusted using the Bayesian shrinkage (sh_log2FC).

### Single cell RNA sequencing analysis

HNEC differentiated in 6-well plastic plates (Cat#657160, Greiner; n=3 independent healthy donors) were dissociated from the culture plate using TrypLE express enzyme and resuspended to a single cell suspension. Samples were pelleted, washed with PBS, resuspended in FACS buffer (PBS0, 1% FBS, 0.5 mM EDTA and DAPI) and strained (35 µm). 376 cells per donor were immediately sorted into 384-well cell-capture plates containing ERCC spike-ins (Agilent), RT primers and dNTP (Promega) using a BD FACSJazz (BD Biosciences). ScRNA-seq was performed according to the SORT-seq protocol ^19^. In short, cells were lysed for 5 min at 65 °C, RT and second-strand mixes were dispensed by the Nanodrop II liquid handling platform (GC Biotech) and double-stranded cDNAs of single-cell was pooled and transcribed following the CEL-seq2 protocol (Hashimshony et al., 2016). Samples were sequenced with 75.000 reads per cell on a Illumina NextSeq. For the analysis of the scRNA-seq data, paired-end reads were aligned to the human transcriptome using Burrows-Wheeler Alignment tool ^39^. Read 1 was used to assign reads to map reads to the correct cells and read 2 was mapped to gene models. Only uniquely mapped reads were used for further analysis. Reads duplicated were removed by excluding reads with identical library, cellular and molecular barcodes. Transcript counts were adjusted to the number of expected molecule based on counts, possible UMI’s and Poissonian counting statistics. In total, 181 cells had to be excluded during quality control measures, leaving 335 cells for donor 1, 284 cells for donor 2 and 328 cells for donor 3. Clustering and analysis of the sequencing results were performed using the Seurat pipeline and BBrowserX (BioTuring). Plots were created with Vinci software (BioTuring).

### High speed video microscopy and ciliary beat frequency analysis

Submerged-differentiated HNEC, cultured in plastic 96-well plates and ALI-cultures in 24-well transwell cultures, were imaged on a Thunder Imager 3D live Cell with a DFC9000 GCT camera (Leica) in a 37°C heated chamber. The presence of cilia in the analyzed videos was confirmed by a life-cilia staining (200 nM SiR-tubulin (Cat#SC002, Spirochrome AG), which was incubated for 4 hours prior to imaging. Cultures were washed twice with PBS prior to imaging and acquisition took place within 30 minutes after washing. Videos were taken with a 40× objective at three randomly picked locations per culture with an imaging speed of at least 200 frames per second (fps).The ciliary beat frequency was estimated using “Temporal ICS” command of Correlescence v.0.0.6 plugin for ImageJ. The full version of the corresponding code is available online (https://github.com/ekatrukha/Correlescence), but in short, it consists of the following steps. At the initial stage, the average intensity image was subtracted from recorded timelapse to remove a static component. Then for each pixel position of the time image stack we calculated normalized autocorrelation function over different time delays. The period of pixel’s intensity oscillations was estimated as a position of a first maximum of the autocorrelation function with a tolerance above 0.2. The frequency was calculated as the period’s reciprocal value. As an output, we obtained an output image of the same X,Y dimensions with its “intensity” values equal to the frequency. All frequencies below 0.1 Hz and above 28 Hz were excluded. To filter for background noise we removed pixel clusters smaller than 25 pixels. For figures and statistics we used the mean CBF value per video.

### Generation of organoid from S-diff HNEC and assessment of forskolin induced swelling (FIS)

Conversion of submerged-differentiated airway epithelia into organoids was conducted essentially as previously described with ALI-differentiated cultures ^16,17^. First, BCs were differentiated at submerged conditions on PureCol coated thermo-reactive Nunc dishes (35 mm) with UpCell Surface (Thermo Fisher Scientific) for at least 18 days. To detach the differentiated epithelial monolayer, the culture medium was substituted for ice-cold Ad-DF and culture dishes were placed on ice for a period of 5-10 minutes. Subsequently, the detached epithelial monolayer was disrupted into epithelia fragments, which were subsequently embedded in 30 µl droplets of basement membrane extract (BME; Cat#3532-010-02, Trevigen). Next, solidified BME droplets were overlayed with airway organoid medium consisting of 95.5% (v/v) Ad-DF with 2% (v/v) B-27 Supplement, serum free, 1% (v/v) GlutaMAX, 10 mM HEPES, 1.25 mM N-Acetyl-L-cysteine, 5 mM Nicotinamide (Cat#N0636, Sigma-Aldrich), 500 μM A83-01 and 1% (v/v) penicillin/streptomycin supplemented with DAPT (5 µM), FGF7 (5 ng/mL) and FGF10 (10 ng/mL). Epithelial fragments, self-organized into organoids (1-2 days), were subsequently transferred in 4 µl droplets of BME on a prewarmed 96-well plate. CFTR function and CFTR modulator responses were determined in a forskolin-induced swelling (FIS) assay as previously described ^17^. CF airway organoids were pre-treated with CFTR correctors: 5 µM VX-809 (Cat#S1565, Selleck Chemicals), 5 µM VX-661 (Cat#S7059, Selleck Chemicals), 5µM VX-445 (Cat#HY-111772, MedChemExpress), or vehicle control for 48 h. CFTR-dependent organoid swelling was measured after stimulation with forskolin (5 µM, Sigma-Aldrich, Cat#F3917-10mg) and the CFTR potentiators VX-770 (5 µM, Selleck Chemicals, Cat#S1144).

### RSV infections in submerged-differentiated airway epithelia

For the infection experiments we used mKate-RSV-A2 for the optimization procedures and RSV-A and -B clinical isolate strains obtained by the INFORM study ^40^. RSV viral strains were propagated in Human Epithelioma-2 (HEp-2) (ATCC CCL-23) cells as previously described ^10^. Briefly, HEp-2 cells were seeded at 2 x 10 ^ 6 cells in a T25 flask one day prior to infection or until 90% of a confluent monolayer was obtained. When confluent, cells were infected with RSV (multiplicity of infection (MOI) = 0.1) and incubated for 3-5 days till 60% of cytopathic effects (CPE) was observed. Virus stocks were snap frozen on dry ice and stored at −80 °C until further use. Viral titer was calculated according to 50% tissue culture infectious dose (TCID50). S-diff HNEC cultured in a 96-wells format and differentiated for 28 days were infected in duplicates. Cell cultures were initially washed once with Dulbecco’s modified Eagle’s medium (DMEM; 12634-028, Thermo Fisher Scientific) at 37 °C, with 5 % CO_2_ for 10 min and subsequently, washing medium was removed and cells were incubated with viral inoculums diluted in 1:1 DMEM : Opti-mem medium at 37 °C, with 5 % CO_2_ for 2 h. When experiments involved treatment application, virus was incubated with Nirsevimab prior to cell incubation for 1 hour. After infection was carried out, inoculum was aspirated and cells were rinsed twice with PBS and left in culture for the defined period of infection upon addition of differentiated medium. Cells were refreshed every two days and negative controls consisted of mock infections, where medium alone was used without virus. To investigate RSV growth kinetics, S-diff HNEC and HEp-2 cells were cultured on 24-wells plate and infected with RSV at a MOI of 0.01 or 3. After 2h of incubation (37 °C, 5% CO_2_), viral inoculum was collected for titration by TCID50 to verify virus input and mentioned as the zero hours timepoint for growth curves. Subsequently, cells were rinsed twice with PBS and differentiation medium was added. For the multi-step growth kinetic assay, at 2, 24, 48 and 72 hours post-infection (HPI), infected cells were scraped in medium and freeze-thawed to release cell-bound virus. For the single-step growth kinetic assay, virus was harvested at 2, 6, 24, 32 and 48 HPI. Viral supernatants of different timepoints were collected and titrated by TCDI50 in HEp-2 cells.

### Cell-based RSV detection ELISA

An indirect enzyme-linked immunosorbent assay (ELISA) was used to quantify RSV F -protein. Cells were fixated prior to staining with 80% acetone in PBS (v/v) at 4 °C for 15 – 30 min. RSV F-protein was detected using a mouse anti-RSV antibody used 1:5000 in casein (MAB8262, Merck), which was incubated at 37 °C, 5% CO_2_ for 1 hour. After incubation, cells were washed 4 times with 0.1% PBS/Tween-20 (PBS/T) and subsequentially, horseradish peroxidase (HRP)-conjugated goat anti-mouse diluted 1:2000 in PBS was incubated at 37 °C, 5% CO_2_ for 1 hour. After a 6-times final washing step with 0.1 % PBS/T, 100 µl/well of substrate solution tetramethylbenzidine (TMB) was added followed by incubation in the dark at room temperature for 7 minutes. Hereafter, 50 µl/well of stop solution (2N H2SO4) was added and RSV specific F-protein levels were quantified by measuring the optical density (OD) per well at 450 nm. Uninfected controls were included on each plate to correct for background signal. The ELISA was validated in the presence of mkate RSV; MOI = 2 or the absence of infection (mock condition), resulting in maximum (max) signal values and minimum (min) signal values. Viral infection was quantified as previously described by measuring OD signal at 450 nm. CV values were calculated according to the following formula: % CV = (sd of means)/ (mean of means) × 100. Max and min infectivity enabled Z’-factor calculation of each 96-well plate according to the following formula: Z’-factor = 1 − (3× (σp + σn)/ (μp − μn)), where σp is the standard deviation of the max signal wells (n = 36 per plate, mkate RSV; MOI = 1), σn is the standard deviation of the min signal wells (n = 36 per plate, mock condition), μp is the mean of the max signal wells and μn is the mean of the min signal wells. Three biological replicates were performed.

### Statistical analysis

GraphPad Prism 9.3.0 was used to perform statistical analyses. Statistical tests and significance are indicated in figure legends.

