## Supplementary information for "Small molecule-directed differentiation of submerged-cultured human nasal airway epithelia for respiratory disease modelling"

Supplementary Fig: 1–5

Supplemental Videos: S1-6

Table S1: Enriched genes of cell type specific clusters in submerged-differentiated cultures based on scRNA-seq

Table S2: Differentially expressed genes submerged-differentiated cultures, ALI-cultures and basal cells based on bulk RNA-seq.

Table S3: Antibodies

Table S4: Primer sequences

### Supplementary figure legends

**Figure S1: The effect of BMP on submerged differentiation and secretory cell characterization.**

a) Representative immunofluorescent images of a HNEC differentiated submerged with and without the Notch inhibitor DAPT, the BMP inhibitor Noggin, or recombinant BMP4 for 18 days. Cells were stained for the secretory cell marker MUC5AC (purple), ciliated cell marker β-tubulin IV (β-tub IV; yellow) and DAPI. b) Quantification of β-tubulin IV (β-tub IV) and MUC5AC signal (n=3 images for 3 independent donors). c) Representative immunofluorescent images of S-diff HNEC stained for the secretory cell markers SLPI, pIgR, and CC10 (purple), together with tubulin IV (β-tub IV; yellow). Data are presented as mean ± SD with individual data point. Statistical significance was tested using a two-tailed paired t-test. ****: $p<0.0001$.

**Figure S2: Bulk RNAseq comparison between S-diff HNEC, BCs, and ALI-diff HNEC.**

a) Representative immunofluorescent images of ALI-cultures differentiated with DAPT and DMH-1 for 18 days. Cells were stained for the secretory cell marker MUC5AC (purple), ciliated-cell marker β-tubulin IV (β-tub; yellow) and DAPI. b) Heatmap showing expression of all DEGs in submerged-differentiated HNEC compared to basal cells (BCs) and ALI-differentiated cultures (n=9 independent donors). c) Normalized mRNA read counts of a selection of epithelial markers in BCs, S-diff, and ALI-differentiated HNEC, including the basal cell markers: *KRT5*, *ITGA6, NGFR, TP63*, and *KRT14*; Secretory cell markers: *MUC5AC*, *SLPI, PIGR, LYPD2,* and *WFDC2*; Ciliated cell markers: *FOXJ1, DNAH5, CDHR3, TUBB4B,* and *CAPS*. d) Heatmap showing marker gene expression of top 20 enhanced and reduced DEGs in S-diff HNEC compared to ALI-HNEC (n=9 independent donors). Data are presented as mean ± SD, and individual data point. Statistical significance was tested using a two-way ANOVA with Dunnett's multiple comparison test *: $p<0.05$, **: $p<0.01$, ***: $p<0.001$, ****: $p<0.0001$.

**Figure S3: Changes in cell composition during submerged differentiation.**

a) Graphic illustration showing the time course experiment set-up. Submerged cultures were used for experiments at six time points between day 0 and day 42 of differentiation. b) Quantitative PCR comparing the expression of *TP63, MUC5AC*, *SPDEF, FOXJ1, CFTR* and *FOXI1* of submerged cultures differentiated for 0, 7, 14, 21, 28, and 42 days (n=2 replicates for 3 independent donors). Statistical significance was tested using a two-way ANOVA with Dunnett's multiple comparison test compared to day 0. c) Representative immunofluorescent images of submerged cultures differentiated for 0, 7, 14, 21, 28, and 42 days. Cells were stained for the secretory cell marker MUC5AC (purple), ciliated cell marker β-tubulin IV (yellow), and DAPI (cyan). d) Quantification of β-tubulin IV and MUC5AC (n= 3 independent donors and n=3 different locations). Data are presented as mean ± SD with individual data point. Statistical significance was tested using a two-way ANOVA with Dunnett's multiple comparison test to day 0. Only significant differences are shown. *: $p<0.05$, **: $p<0.01$, ***: $p<0.001$, ****: $p<0.0001$.

**Figure S4: Long-term expansion and submerged differentiation of S-diff HNEC.**

a) CBF in Hz of submerged cultures differentiated for 0, 7, 14, 21, 28, and 42 days (n=3 independent healthy donors). Each point represents the mean frequency for one video. b) Graphic illustration showing the long-term expansion of BCs in 6-well plates and differentiation in 96-well culture plates at passage seven (p7) and p13. c) Population doublings (PD) measured in BC cultures from donors HC (5,6,10) and PCD1-5. PD=3.32×(log(cells harvested/cells seeded)). d) Representative brightfield images of BC cultures from a HC and PCD subject at p6 and p12. Images were taken before passaging. e) Quantification of β-tubulin IV and MUC5AC signal measured in p7 and p13 S-diff HNEC of HC (n=3 independent subject) and PCD subjects (n=4 independent subjects). f) CBF in Hz of submerged-differentiated HC (n=3) and PCD-donor derived (n=5) cultures at p7 and p13. Each point represents the mean frequency for one video. Data are presented as mean ± SD, and individual datapoints. Statistical significance was tested using (a) a Dunnett's multiple comparison test to day 21. (e,f) a Tukey's multiple comparison test. ns = non-significant, *: $p<0.05$, ***: $p<0.001$, ****: $p<0.0001$.

**Figure S5: Validation experiments of RSV infections.**

a) Live imaging of mKate-RSV-A2 infections with different MOIs of ALI-differentiated HNEC at 3 DPI. b) HEp-2 cells were infected with RSV-A2 mKate at MOI = 0.01 or 3. Virus growth kinetics was determined by titrating supernatant and cell-bound RSV at different time intervals following infection. c) Replicate experiments of three 96-wells plates to assess the reliability and robustness of the ELISA assay in which n=36 wells were infected with RSV-A2 mKate at an MOI of 2 at 3 DPI (=max OD signal) and in which n=36 wells S-diff HNEC remained uninfected (=min OD signal). CV values were calculated according to the following formula: % CV = (sd of means)/ (mean of means) × 100. Z'-factor of each 96-wells plate was calculated according to the following formula: Z'-factor = $1 - (3\times (\sigma p + \sigma n)/ (\mu p - \mu n))$, where σp is the standard deviation of the max signal wells (n=36 per plate, RSV-A2 mKate; MOI = 1), σn is the standard deviation of the min signal wells (n=36 per plate, mock condition), μp is the mean of the max signal wells and μn is the mean of the min signal wells. d) Correlation between qPCR (fold change) and ELISA (OD450nm). e) Cell-based ELISA with S-diff HNEC, VERO E6, and HEp-2 cells, infected with different MOIs of a RSV-B clinical isolates. Data are presented as mean ± SD, from two individual experiments with two technical replicates per condition in each experiment; SD is indicated by error bars.
