## Supplementary figures and images for "Small molecule-directed differentiation of submerged-cultured human nasal airway epithelia for respiratory disease modelling"

### Figure S1

Supplementary Figure 1

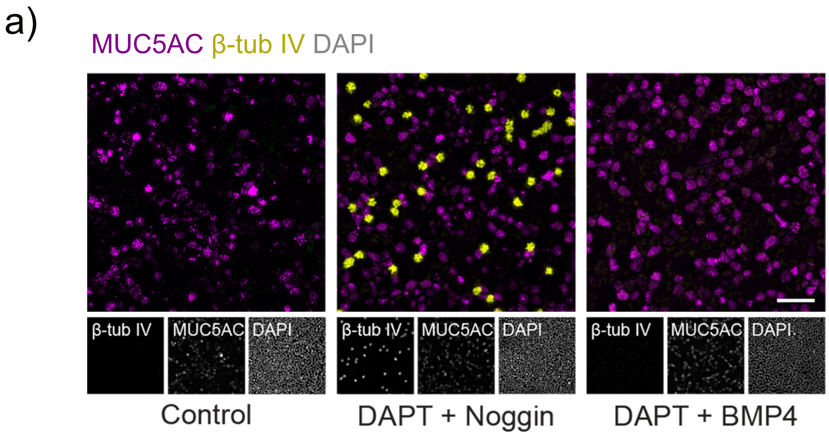


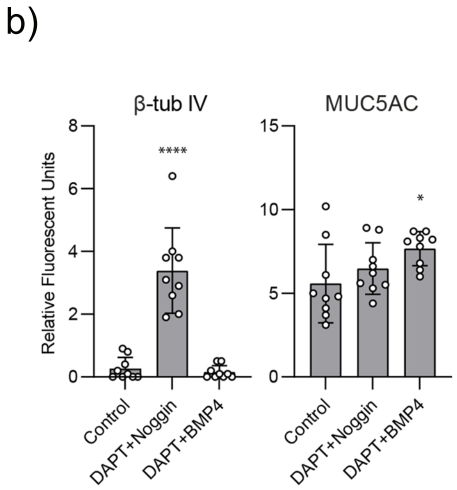


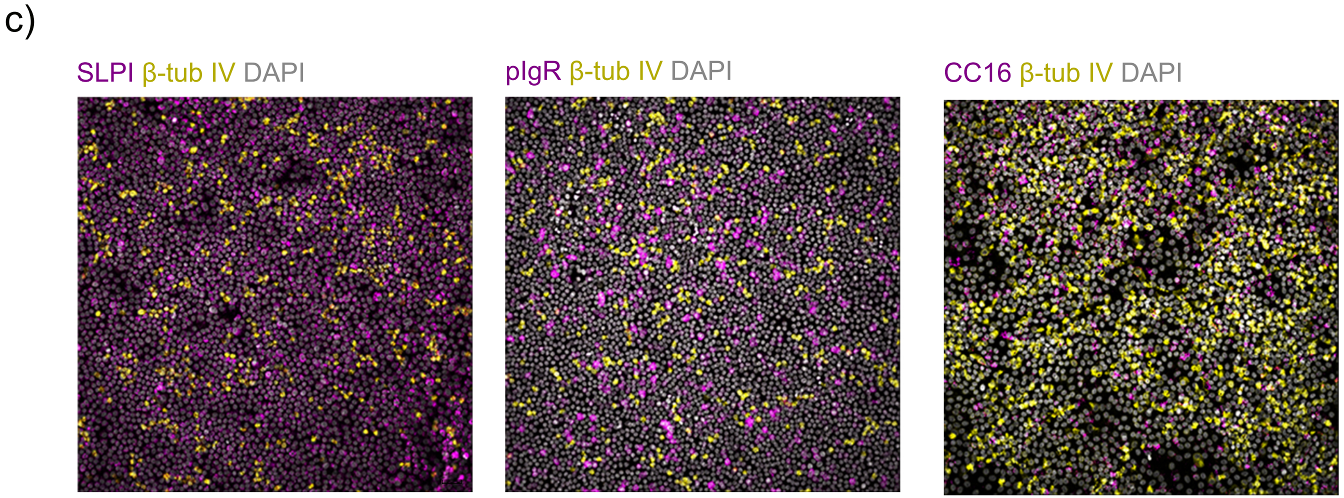

### Figure S2

Supplementary Figure 2

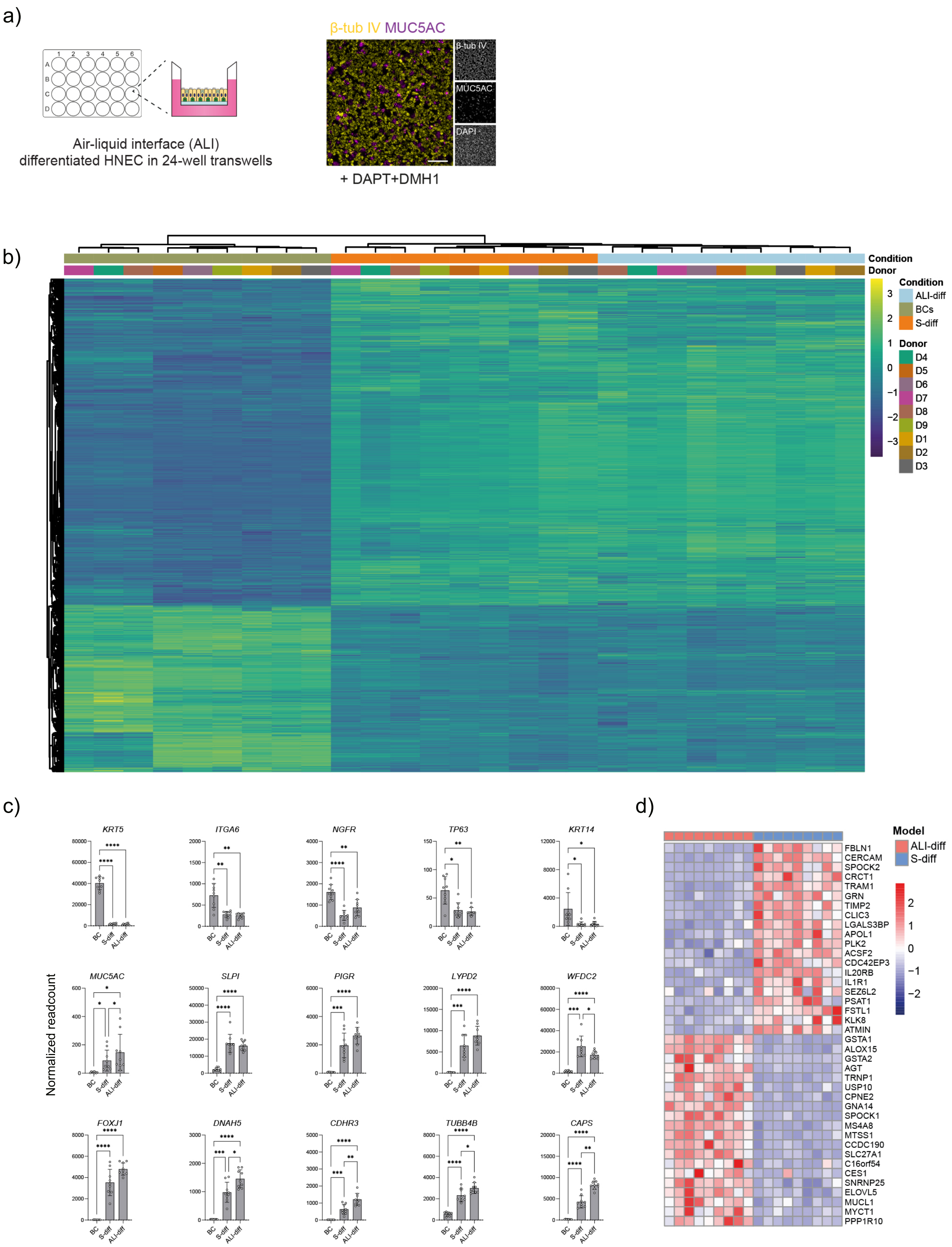

### Figure S3

#### Supplementary Figure 3

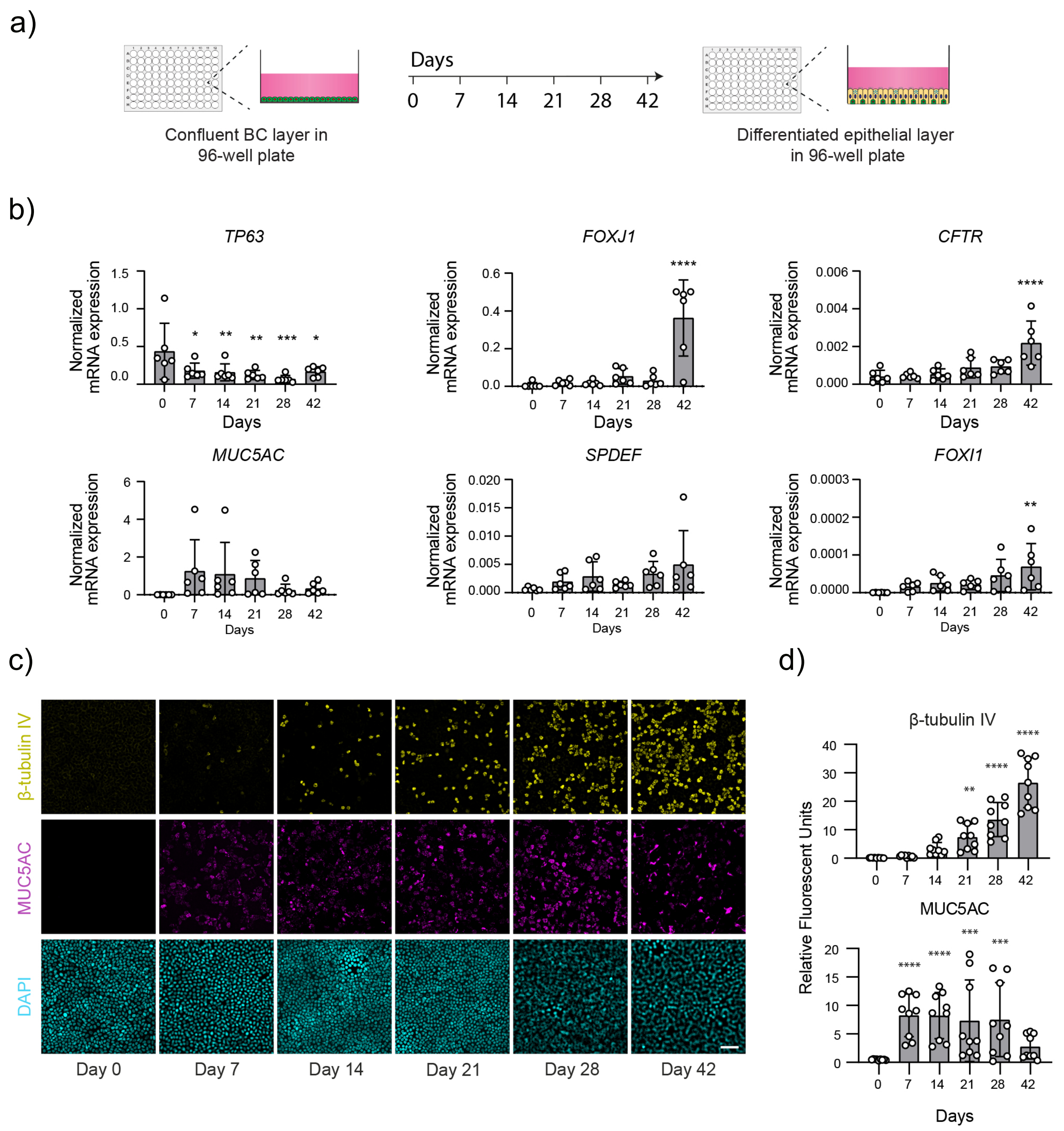

### Figure S4

#### Supplementary Figure 4

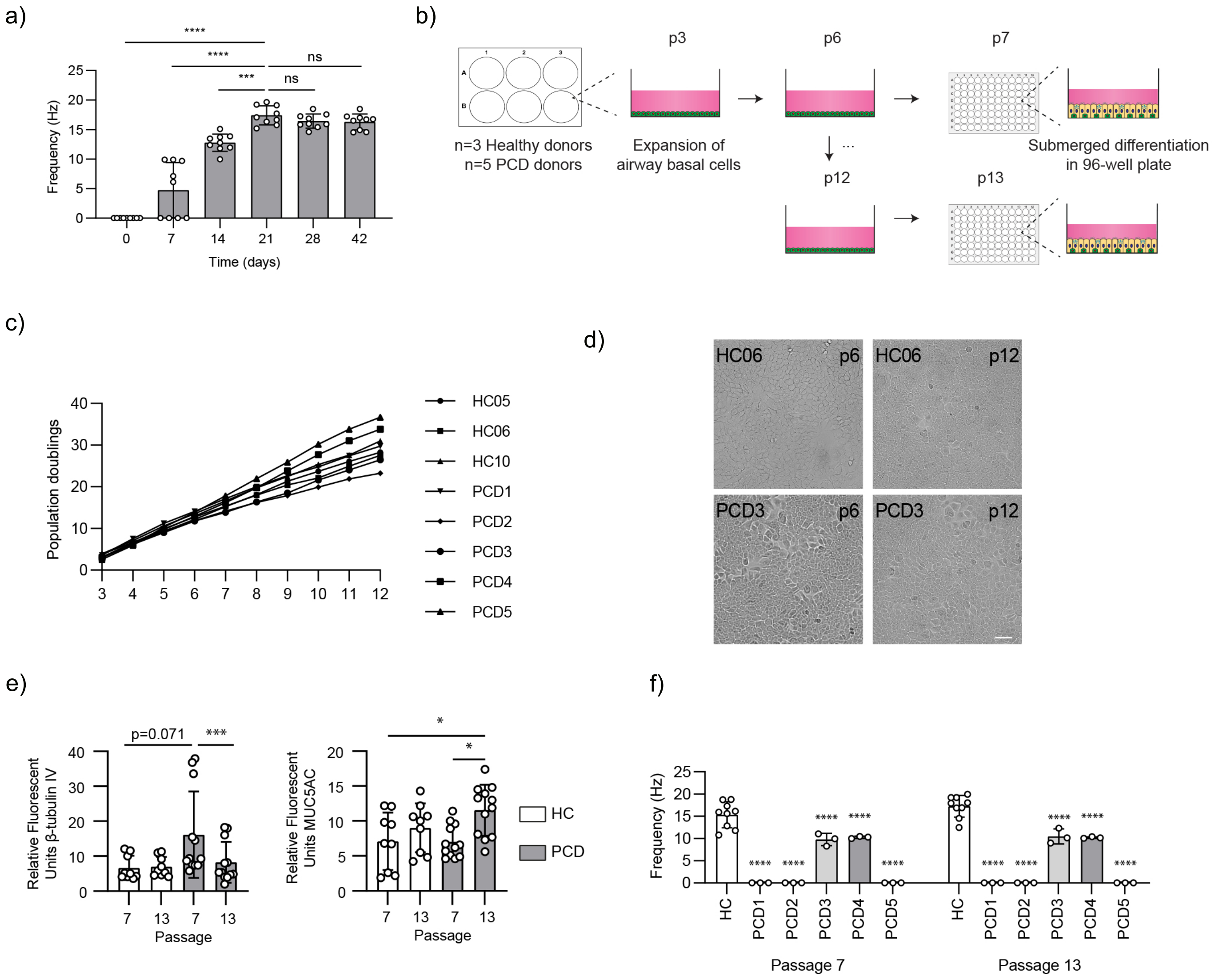

### Figure S5

#### Supplementary Figure 5

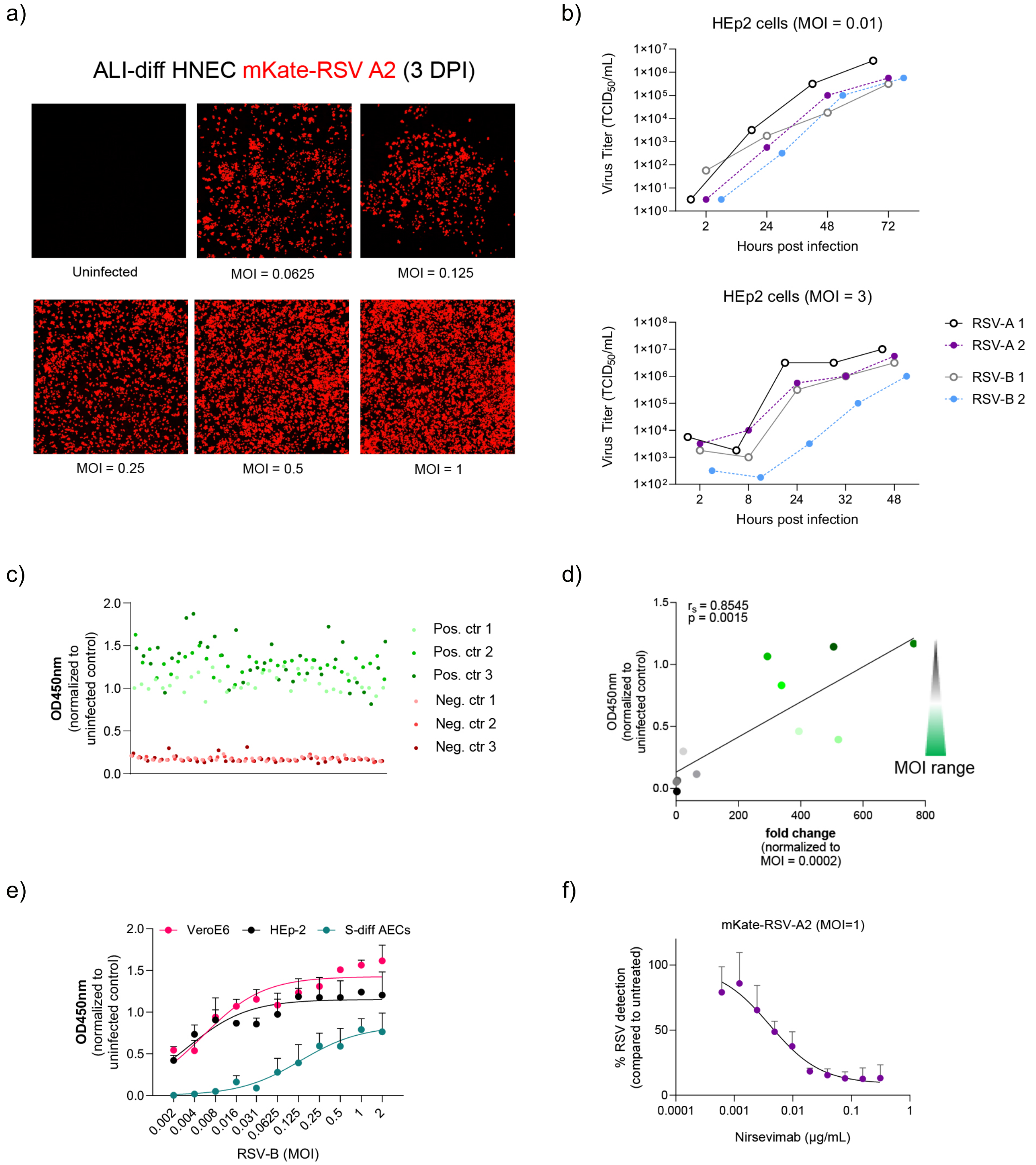
